# Decisional reference point pathology: a mechanism and marker for major depressive disorder in humans

**DOI:** 10.1101/2025.04.18.649538

**Authors:** Aadith Vittala, Lulu Wu, Dongni Yan, David Liebers, Elizabeth Tell, Xiaotong Song, Damon Dashti, Kenway Louie, Candace Raio, Dan V. Iosifescu, Paul Glimcher

## Abstract

The decisional *reference point*, the central mechanism of behavioral economics, conditions our evaluations of reinforcers. It determines whether a given event is experienced as positive or negative. Here we show, for the first time, a significant and pathological elevation of the reference point in patients with depression that correlates with disease severity. We also show that the mechanism for reference point setting is profoundly impaired in patients with depression. These findings link the previously demonstrated treatment of depression by deep brain stimulation to modulation of the reference point in the anterior cingulate cortex and identify pathology in the dynamics of reference point setting as a novel mechanism in the disorder. Finally, these results lay the foundation for a three-minute virtual diagnostic test for depression.

## Introduction

Major depressive disorder (MDD) is a debilitating and common psychiatric illness that impacts 185 million people worldwide (*1*). Despite its prevalence and impact, scientists have not yet identified the cognitive mechanisms that underlie the disorder. Tests for depression and scales for measuring disease severity reflect this lack, relying on catalogs of symptoms rather than specific measures of an underlying pathology (*2*). While depression likely reflects more than one cognitive mechanism (*3*), identifying a core mechanism and a disease-specific pathology of that mechanism would constitute a major advance in our ability to both diagnose and treat depression.

Since Kahneman and Tversky’s landmark theories of decision-making were introduced in the 1970s, it has been widely appreciated that the hedonic quality of a reinforcer, whether it is experienced as positive or negative, depends in part on the decision-maker’s expectations, their reference point (*4*–*10*). For example, a single slice of pizza is positively reinforcing if one expects no food but negatively reinforcing if one expects (either consciously or unconsciously) a five-course dinner. Working from this idea, Rigoli and colleagues (*11, 12*) and Glimcher and Tymula (*13*) have previously hypothesized that depression may reflect a specific pathology of the reference point representation in the brain, leading to a failure to experience positive reinforcers as positive and therefore to depression. Neural data provides some support for this hypothesis. We and other groups have demonstrated the existence of a neural representation of the decisional reference point in the anterior cingulate cortex of monkeys (*14, 15*). This same region appears to be implicated in depression: in humans, brain imaging data has pointed towards depression-specific pathological activity in or near the anterior cingulate (*16*), neural activity in this area can be used to decode depression severity (*17*), and the outputs of this area are also believed to play a role in depression (*18*–*20*). Finally, deep brain stimulation of a pathway connecting this region to the frontal cortex and amygdala can be used to treat depression under at least some conditions (*17, 21*–*23*). However, there has been no evidence that the reference point itself is pathological in depression, nor has a plausible cognitive mechanism for such a pathology been suggested.

To test the hypothesis that depression results from a pathology of the decisional reference point and to provide a mechanism by which that pathology might arise, we conducted two experiments in patients diagnosed with major depressive disorder (n=50) and in healthy controls (n=70). The first deployed a validated technique for measuring an individual’s current reference point in a video game-like task modeled on the marginal value theorem from ecology (foraging task) (*24, 25*). In this task subjects forage for apples in an orchard in a manner that allows us to rapidly and accurately measure an individual’s situation-specific decisional reference point. The second was designed to begin to test the hypothesis that the cognitive mechanism which sets the situation-specific reference point may be the source of this pathology. To assess the dynamical process by which the reference point adapts to a given situation or environment, we used a validated snack food bidding task (bidding task) (*26*), modeled on visual adaptation experiments that assess perceptual reference points (*27*) and their temporal evolution after transitioning to a new environment.

Overall, we found evidence that patients with MDD exhibited a remarkably elevated situation-specific reference point. In the foraging task, we found that the static reference point in patients with major depressive disorder (across all levels of disease severity) was on average 50% higher than that of healthy controls. We found that the specific degree of elevation of the reference point was highly correlated (r = 0.71) with depression severity, as would be expected if this was a core mechanism of the disorder. Finally, we found that with just a few minutes of task data we could separate patients from controls with very high discrimination (an area under the receiver operating curve of 0.85).

Our second, and perhaps more important finding, was that patients showed a pathological inflexibility in how they adapt their reference points to dynamically changing environmental quality – a mechanistic dysfunction which might underlie the disorder. While healthy controls flexibly adapt their reference points to the current environment using a largely linear and exponentially relaxing process, patients showed a highly distinctive nonlinear and non-exponential mechanism for reference point setting.

These findings provide significant support for the existing hypothesis that major depressive disorder is associated with a pathology of the decisional reference point. More importantly, our study of reference point dynamics in patients may offer the first identification of a pathological cognitive mechanism that may underlie the disorder. Irrespective of underlying mechanism, our findings also identify the foraging task as a low-cost, high-reliability, mobile-friendly tool for remotely assessing depression severity in as little as three minutes. Smartphone or web-based delivery of such a tool could speed the process of treatment selection, reducing the burden of depression for patients in the future.

## Results

### Involvement of the Reference Point in MDD: Foraging task

To assess our hypothesis that a reference point pathology may be a significant and previously undocumented feature of major depressive disorder (Figure 1a), we recruited participants with a current clinically verified diagnosis of depression (n=50; participant characteristics in Table S1) and healthy controls (n=70) to complete two tasks to assess their reference point: a foraging task (*25*) (Figure 1b) and a bidding task (*26*) (Figure 1c). In the foraging task, participants were instructed to collect as many apples as they could from a series of cartoon apple trees. At the end of the experimental session participants were compensated $0.01 for each apple harvested over the entire session. On each trial, a participant could choose either to harvest apples from the current tree, or to travel to a new (unharvested) tree to begin harvesting there. Each time a given tree was harvested, it yielded a sequentially diminishing number of apples. Subjects had to decide when the diminishing number of apples was no longer positively reinforcing, at which point they should opt to travel to a new (unharvested) tree. In a given block of trials, new trees were either 3s (short-travel block) or 10s (long-travel block) away. Participants completed four alternating 7-minute blocks (counterbalanced for order across subjects). The optimal leaving criteria (the number of apples which signals that staying to harvest again diminishes one’s total reward rate) is defined under these conditions by the marginal value theorem (*24, 25*), and it also conforms to the standard definition of the reference point from behavioral economics (less the secondary consumption utility component) (*8*).

**Figure 1.**
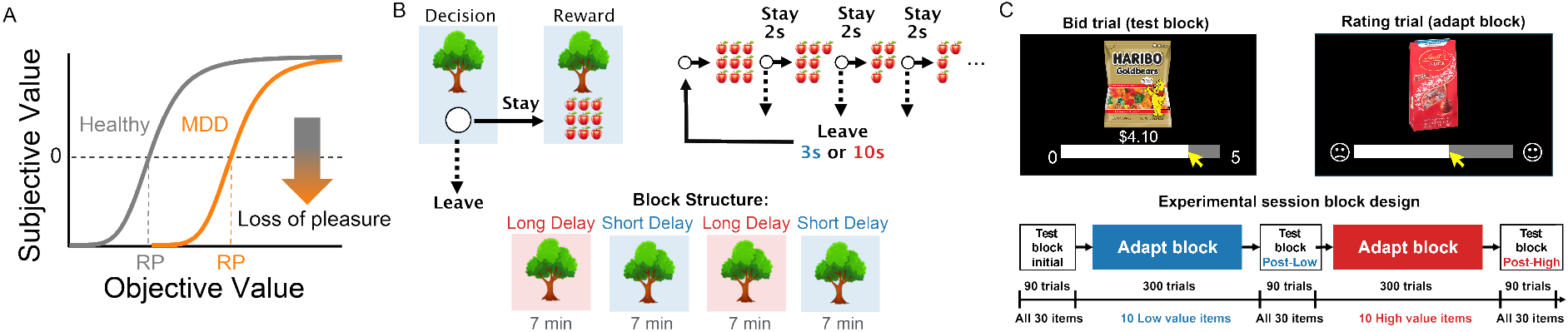
Hypothesis and procedures for foraging (static reference point) and bidding (reference point dynamics) tasks. A) We hypothesize that depression (MDD, orange) is associated with an increased decisional reference point (RP) compared to controls (gray), leading to reduced subjective valuation across a wide range of reinforcers B) Foraging task. Participants completed a sequential decision-making task where they chose either to continue harvesting apples from a gradually depleting tree, or to travel to a new un-depleted tree and begin harvesting there. Within a block of fixed 7-minute duration, choosing to leave imposed either a 3s (short) or 10s (long) delay. Participants completed four blocks presented in the order long-short-long-short or short-long-short-long. C) Bidding task. Participants first reported the most that they would pay (from a $5 study endowment) for each of 30 snack food items using an incentive-compatible procedure (baseline). They then experienced adaptation blocks in which only high value or low value snacks were presented. After adaptation, subjects again reported the maximum value they would pay for the original 30 snacks over 90 trials. Systematic changes in bid values reflect the impact of adaptation on the reference point.

As shown in Figure 2a, an internal subjective value curve relates the objective number of apples received on a given trial to the subjective level of reinforcement it produces. We hypothesize that patients with major depressive disorder experience the transition from positive to negative reinforcement (i.e. the reference point) at a higher number of apples than do healthy controls. Figure 2b plots the actual number of apples that triggered each of two participants to stop harvesting and leave for a new tree over a representative 7-minute block. Note that the patient tends to leave after their harvest diminishes to about 8 apples (with significant tree-to-tree variation) while the healthy control tends to depart after rewards diminish to 4 apples. We found that this was consistently true. Across both long- and short-travel blocks, participants with major depressive disorder had a consistently higher average reference point than did healthy controls (Figure 2c, two-sample t-test, short-travel block: p = 2e-8, long-travel block: p=2e-10). This difference did not result from a reduced ability to obtain rewards overall. On average, patients achieved the same long-run rate of reward as did healthy controls (Figure S1: long-travel and short-travel blocks: two-sample t-test, p>0.05).

**Figure 2.**
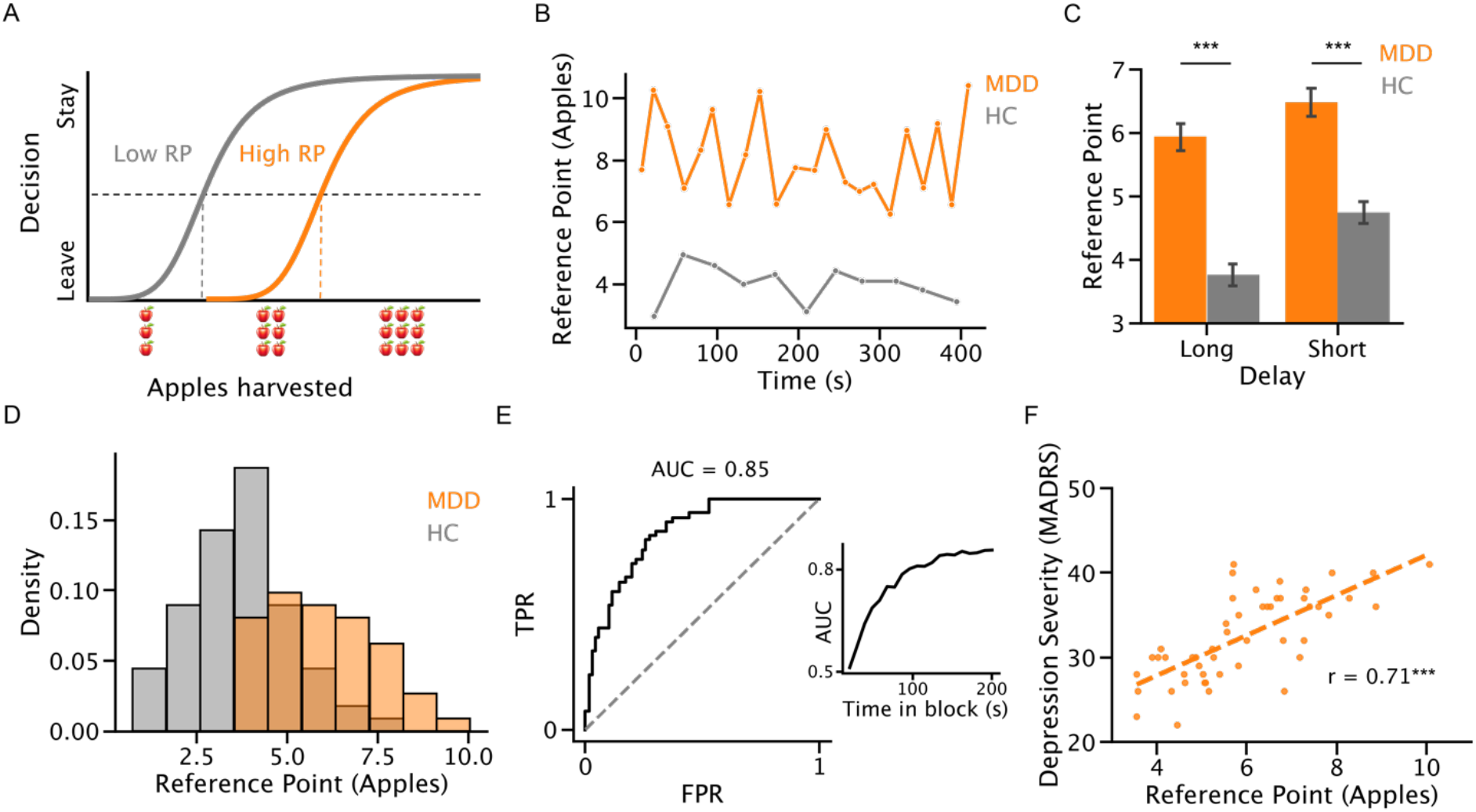
Foraging task: Depressed patients (n=50) showed a higher reference point than healthy controls (n=70) and individual reference points in patients correlate with depressive severity. A) The foraging task measures the static reference point as the threshold number of apples at which participants leave for a new tree. Vertical dotted lines show the border between positive and negative reinforcement for patients (orange) versus healthy controls (gray). B) Observed number of apples in a harvest that triggered the subject to leave for a new tree within a full block of trials (long-travel time condition) for a representative patient (orange) and healthy control (grey). C) Average number of apples that triggers leaving, across all blocks and participants. Error bars are 95% confidence intervals. Differences between the two populations are significant: long-travel block: two-sample t-test, p = 2e-10. Short-travel block: two-sample t-test, p = 2e-8. D) Distribution of average long-travel block leaving points for individual patients and healthy controls. Differences between the two distributions are significant: Kolmogorov-Smirnov test, p=3e-9. E) Area Under the Curve analysis (AUC) using average long-travel block leaving point to predict whether participants are control or MDD. AUC is significantly different than random: Bootstrap N=2000, p<5e-4. Inset: AUC as a function of measurement duration produced by sampling the initial several minutes within the first long-travel block. AUC asymptotes within about 3 minutes. F) Individual average leaving times among patients (long-travel blocks shown) correlate with symptom severity by MADRS score. Linear regression: r = 0.71, p = 6e-9.

When examining individual participants, we found that the average reference point was distributed differently between patients and control participants (Figure 2d), with the distribution of average leaving times being significantly shifted to the right in patients compared to controls (Kolmogorov-Smirnov test, p=3e-9). Employing a standard receiver operating characteristic analysis (*28*) to estimate the criterion-free categorical accuracy of the behaviorally measured reference point in identifying depression, we find that the long-travel blocks yield an area under the curve (AUC) of 0.85 (Figure 2e; similar AUC for short-travel block, see Figure S2). Reducing sample time has no significant effect on AUC down to a measurement time of 3 minutes (Figure 2e inset), suggesting that a clinical assessment of depression with an AUC of ∼0.85 could be achieved in as little as 3 minutes. Perhaps even more strikingly, at the individual patient level, the average reference point we observed for each patient strongly correlated with that individual’s depression severity as measured with the MADRS structured interview (Figure 2f; Pearson’s correlation r = 0.71, p = 6e-9). We examined the test-retest reliability of this 7-minute measure of depression severity and found it to be 0.84 (Figure S3), a number comparable to that achieved with structured clinical interviews (*2, 29*).

### Mechanism for Reference Point Pathology: Bidding task

Having established that the reference point itself is pathological in depression, we set out to test the hypothesis that the cognitive mechanism which sets the situation-specific reference point may be the source of this pathology. Employing the same participants recruited for the foraging task, we asked these individuals to complete the bidding task (*26*) which has previously been shown to assess reference point dynamics in healthy control subjects (*26*). In a baseline block, participants indicated the most that they would pay for each of thirty snack food items presented in a randomized order three times (using the incentive compatible BDM mechanism (*30*)). To dynamically adapt the reference point to a higher (or lower) value, we next presented each individual with their top ten (or bottom ten) snack foods and asked them to rate the desirability of those items for 300 trials. This change in environment elicits a compensatory shift in the reference point in healthy individuals (Figure 3a). After this adaptation block, participants were then asked to again report the maximum they would pay for each of the thirty original objects, three times, to assess the dynamics of the behaviorally measured reference point as it readapted to the original environment. High and low adaptation blocks were presented in sequence to each participant with order counterbalanced across participants.

**Figure 3.**
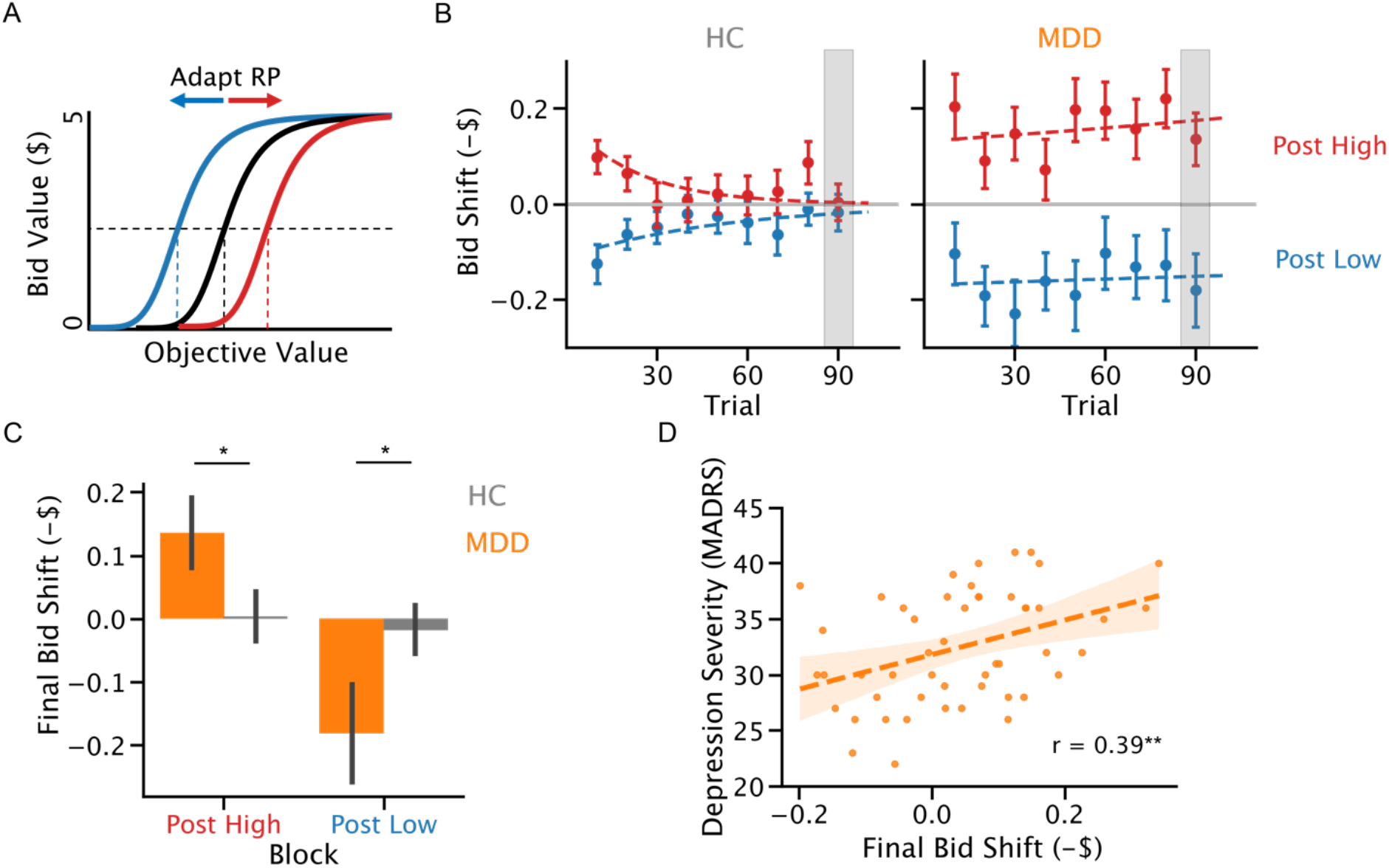
Bidding Task: Depressed patients (n=50) show a reduced ability to meaningfully adapt their reference points to a changing environment compared to healthy controls (n=70). A) Bidding task allows measurement of reference point dynamics. Pre-adaptation to low value items (blue) should transiently shift the reference point to the left and briefly lead to lower bid values for all items. Pre-adaptation to high value items (red) should increase reference point and briefly lead to higher bid values for all items. As subjects adapt to the standard measurement environment these pre-adapted changes should gradually relax back to preadaptation levels. Note that reference point moves opposite to direction of bid shifts. B) Average bid shifts (in negative dollars, over 90 bids/5 min, error bars are 95% confidence intervals) return back to baseline after exposure to a better (high, red) or worse (low, blue) environment. Unlike healthy controls, patients show no significant relaxation of their bids back to preadaptation levels during the 90 trial (5m) measurement interval. Exponential decay (dashed lines) fits well to data from healthy controls but not to data from patients (controls: bootstrapped N=2000 on decay parameter, p=0.01 post-low and p=0.05 post-high, MDD: bootstrapped N=2000 on decay parameter, p > 0.05), C) Residual reference point shift observed during the last ten trials of a measurement block (gray area in panel B) is significantly different between patients and healthy controls. Mean residual shift is shown, and error bars are 95% confidence intervals. Post-high: two-sample t-test, p=0.02. Post-low: two-sample t-test, p=0.02. D) The magnitude of this residual shift after 80 measurements (post-high adaptation shown) is correlated with depression severity in patients: Linear regression: r=0.39, p=0.005.

After adaptation to the high (low) value environment, healthy participants underbid (overbid) briefly, but rapidly readapted to the standard environment (Figure 3b). In contrast, patients showed no detectable readaptation to the standard environment over the 90-trial bidding block. Exponential fits to the bid deviation induced by the adaptation of the reference point show a significant decay back to baseline for healthy controls but not for patients (bootstrapped N=2000, p < 0.05 for controls, p > 0.05 for patients), a striking difference in how patients adapt to their reward environment. Importantly, despite the difference in dynamics, patients still shift their bids both up and down in response to the adaptation block, and their initial shift is the same size as healthy controls (Figure S4, two-sample t-test p > 0.05).

To quantify this change in the underlying adaptational mechanism, we measured for each participant the bid shift from baseline during the last 10 trials of each bidding block. Figure 3c plots the magnitude of this residual adaptation at the end of the blocks. Patients showed little or no readaptation – a failure to adapt to the reward environment – under both conditions (two-sample t-test, p = 0.02). Finally, we found that the magnitude of this failure to readapt normally during the 90-trial bidding block, as measured by residual adaptation in the last ten trials, significantly correlated with depression severity in the patients (Figure 3d; Pearson’s correlation r=0.39, p=0.005).

## Discussion

We demonstrate here that two behavioral assays of the human decisional reference point can be used to discriminate between healthy controls and those with major depressive disorder. Using Constantino and Daw’s foraging task (*25*), we found that patients with major depressive disorder had significantly elevated reference points relative to healthy controls. The foraging task measures the reward level (in apples) at which a participant shifts from finding a tree positively reinforcing (and hence continues harvesting) to finding the tree negatively reinforcing (and hence stops harvesting). For patients with major depressive disorder, that boundary was shifted upwards by about 50% relative to healthy controls. While healthy controls typically found 6 apples positively reinforcing, on average patients found that same number of apples *negatively* reinforcing. Indeed, if we use classical foraging theory to estimate the optimal positive/negative reinforcement boundary (or reference point) which yields the maximal harvest, patients with depression are observed to abandon trees as negatively reinforcing long before reaching that point. Perhaps even more significantly, we find that individual reference points correlated with disease severity as assessed by the well-established Montgomery-Asberg Depression Rating Scale (MADRS), as would be expected if this was a core feature of the disorder. At a more practical level, we note that using our approach measurement of the reference point can be made on a computer or smartphone in as little as 3 minutes, with test-retest reliability similar to that of the gold-standard in-person clinical assessment, the MADRS (*2, 29*).

We also found evidence for pathology in the cognitive mechanism that sets the reference point into alignment with the current environment or situation. The dynamics with which the reference point responds to changes in the environment showed a clear pathology in patients with major depressive disorder. In both foraging theory (*24, 31*) and decision science (*4, 8*) it has been noted that the reference point should reflect a rational expectation about the average value of the environment. We have previously shown that when healthy subjects are asked to rate how much they desire a snack food presented for 300 trials in high or low valued environments, they shift how much they are willing to pay for all snack foods in the original testing environment (*26*).

This shift, however, readapts rapidly to baseline when they are returned to the original environment in healthy controls (*26, 27, 32*), returning to baseline levels with a time constant of roughly 20 trials (∼2 minutes) in this task. Here we found that while exposure to 300 rating trials produced normal-sized shifts in the reference point of the patients, those shifts did not adapt back at all during our measurement period of 90 trials, suggesting an unusual and perhaps non-linear impact of the environmental quality on the reference points of patients with major depressive disorder. This inability of patients to adapt their reference point to the current environment stands out as a novel mechanistic explanation for depression that merits future investigation.

Taken together, our two sets of findings present converging evidence for the proposal that a pathology of the human reference point underlies at least some forms of major depressive disorder (*11*) and provides a novel mechanistic explanation of how this pathology might arise. We note that our findings contrast with those of a study that employed a different but similar patch foraging task (*33*) in depressed patients and controls but observed no significant differences. One critical distinction between the tasks was that Bustamante and colleagues examined the trade-offs between physical and cognitive effort during foraging while ours simply examined efficiency in decision-making. This difference suggests an interesting avenue for future therapeutic research: whether effort can be used to more effectively regulate the reference point in patients with pathological reference point dynamics in more passive decisional or evaluative settings. Our findings, of course, also relate to a more general body of literature examining reward processing deficits in depression. The reference point is known to affect all facets of reward processing, and pathology in the reference point may underlie the impairments in reward bias, option valuation, and reinforcement learning that have been observed in depression (*20, 34*–*37*).

### Relation to existing Neurobiological Data

A number of studies have suggested that in healthy non-human primates, the neural representation of the reference point can be found in the anterior cingulate cortex (*14, 15*). Recent electrophysiological work in human patients has shown that depression severity can be decoded, using machine learning techniques, from activity patterns in a homologous area in humans (*17*). Both of these findings appear to corroborate the conclusion that some forms of depression may reflect a specific pathology of the human decisional reference point as instantiated in the anterior cingulate cortex. Mayberg and colleagues’ observation that electrophysiological modification of the output of the far anterior (subgenual) cingulate cortex is an effective treatment for depression in some patients appears to further strengthen this conclusion (*21*–*23*).

While our findings and those of Mayberg’s group do suggest a specific neurocomputational pathology for some forms of depression, they may also suggest a new avenue for developing treatment strategies – particularly with regard to the altered reference point dynamics in depression. Anhedonia is an often-undertreated symptom of depression associated with negative affective consequences; many patients seek treatment for depression not just normalize mood, but to restore hedonic pleasure (*38, 39*). The theoretical framework of reference point pathology that our results suggest could more broadly lead to the development of better tools for quantifying anhedonia and for the development of new pharmacological and behavioral interventions. Future measurements of the reference point as a biomarker for major depressive disorder in larger and more heterogeneous populations could help stratify patients for treatment as well.

Our findings leave unresolved the question of whether pathological shifts in the reference point are specific to major depressive disorder or represent a transdiagnostic feature of other related disorders associated with anhedonia. Indeed, the single category of depression is increasingly thought to involve several sub-types of the disorder (*3*), which may be heterogenous for reference point pathologies. One fundamental limitation of our study has been the necessity to group subjects with potentially different subtypes of depression together. It may be the case that measuring the reference point will, in the future, identify a specific subtype of depression. If so, by more accurately identifying a selective subset of depression in larger future studies, our measurements might help define the heterogeneity of processes that give rise to depression.

## Methods

### Participants and clinical assessment

Seventy healthy adults (mean age 35, thirty-three female) and fifty adults with depression (mean age 36, twenty-two female) participated in this study. All participants gave informed consent in accordance with the Institutional Review Board of the New York University School of Medicine. In order to be eligible for the study, depressed patients had to meet DSM-5 criteria for major depressive disorder as assessed by the Mini International Neuropsychiatric Interview (MINI) (*40*), have a current Montgomery-Asberg Depression Rating Scale (MADRS) (*29, 41*) score greater than 19 and Beck Depression Inventory (BDI) (*42*) score greater than 19. They also could not have a primary anxiety disorder, a psychotic disorder, or any other mood disorders. Healthy controls were required to have no psychiatric conditions (diagnosed with MINI), and BDI score less than 14 (to exclude sub-syndromal depression). MADRS scores were not obtained for healthy control subjects.

All subjects participated in a single three-hour test session. Participants first completed a demographic information form, an abbreviated version of the Raven’s Progressive Matrices, and self-rated depression scales. Next, depressed patients underwent clinical assessment with a trained interviewer on the MINI, MADRS, and Columbia Suicide Severity Rating Scale^5^.

Finally, all participants completed the foraging task and the bidding task.

### Foraging task

The foraging task is a gamified set of decisions in which participants make serial choices in a virtual patch-foraging task (*25*). On each trial, participants were presented with a fixed image of a tree and had to decide whether to harvest apples from that specific tree or whether to move on to a fresh and unharvested apple tree. Participants indicated their choice with one of two key presses. If they decided to harvest, two seconds elapsed and then the number of apples returned from the tree was displayed. After each sequential harvest from a given tree, the number of apples decayed by a multiplicative factor chosen from a beta distribution with an average decay rate of 12%. The number of apples harvested was discretized to a single apple by rounding to the nearest whole number. In this way, a given tree yielded fewer apples on sequential harvests.

If participants chose to move to the next tree, they incurred a fixed travel delay (either 3s or 10s depending on block type), after which a new animated tree was revealed. New trees provided an initial harvest normally distributed around a mean of 10 apples with a standard deviation of 1 apple. Each block of trials took a total of seven minutes, and to ensure that reaction time did not influence the total number of apples harvested, it was included within the total travel or harvest delay times. Before starting the task, participants completed a training session where they experienced each travel time and acclimated to making harvest-vs-travel decisions. During the task, participants completed four blocks, and each block clearly signaled whether it was a short or long travel time block. For half of all subjects, block order was long-short-long-short with the other subjects seeing a short-long-short-long ordering. At the end of the four blocks, participants were shown their total winnings in number of apples, and this number was converted into dollars such that optimally performing participant could win around $30. On average, participants won $27 ± $3 based on their actual choices. There was no difference in amount won between patients and controls (see Figure S1).

### Bidding task

Participants completed the bidding task as previously described in (*26*). Before the task began subjects were endowed with $5 by the experimenters with which to bid on the presented snack foods. The task consisted of three bidding blocks separated by two adaptation blocks. Each bidding block included 90 total trials in which subjects reported their maximum willingness to pay for 30 individual snack food items familiar to American subjects. In each trial, participants saw an image of the individual item on a computer screen and selected a bid amount between $0 and $5 on a slider bar. Each of the thirty items was presented three times in a randomized order. After all trials were complete, a single randomly selected trial from the three bidding blocks for realization. The bid was realized using a standard Becker-Degroot-Marschak experimental economic process which had been carefully explained to the subjects before bidding began, a procedure designed ensure that participants are incentivized to reveal the maximum amount that they were willing to pay for each snack food item (*30*).

After the initial bidding block was complete, items were ranked by the subject’s mean bids and the ten highest and lowest value items were selected for use in the subsequent adaptation blocks. During the two adaptation blocks, subjects were presented with 300 trials in which they viewed a picture of a snack food item (drawn from either the ten highest or lowest value items for high- and low-adaptation blocks respectively) and rated the “pleasantness” of the item using a visual-analog scale implemented as a slider bar. Immediately following each adaptation block, subjects completed the next bidding block and bid on all 30 items three times each. The presentation order of high- and low-adaptation blocks was randomized across subjects.

### Data analysis

All data analysis and statistics was conducted with custom-written Python scripts. To quantify reference point shifts in the bidding task, data was aggregated across groups and fit to an exponential decay model: y = B0 exp(−lambda t) where the parameters B0 and lambda represent the initial value and decay constant respectively. Data from post low-adaptation and post high-adaptation bidding trials were fit separately to this model. Significance of exponential decay parameter was determined by bootstrap resampling (2000 iterations).

## Data availability

The deidentified data that support the findings of this study are available from the corresponding author.

## Code availability

Scripts to run the analysis are available from the corresponding author.

## Author contributions

AV: study design, data collection, drafting manuscript, statistical analysis. LW: study design, data collection, statistical analysis. DY, ET, XS, DD: data collection. DL: clinical assessment of participants. KL, CR: study design, comments on manuscript. DI: conception and design of study, clinical assessment of participants, comments on manuscript. PG: conception and design of study, comments on manuscript.

## Ethics declaration

In the last 10 years, DVI has served as a consultant for Alkermes, Allergan, Angelini, Autobahn, Axsome, Biogen, Boehringer Ingelheim, the Centers for Psychiatric Excellence, Clexio, Delix, Jazz, Lundbeck, Neumora, Otsuka, Precision Neuroscience, Relmada, Sage Therapeutics, and Sunovion. He has received grant/research support (paid to his institutions) from Alkermes, AstraZeneca, Brainsway, LiteCure, NeoSync, Otsuka, Roche, and Shire. All other authors declare no competing interests.

