## Supplemental Figures for "Decisional reference point pathology: a mechanism and marker for major depressive disorder in humans"

**Supplementary Data**

Table 1. Participant characteristics

| | Group | |
|---|---|---|
| | Healthy controls | MDD patients |
| Number | 70 | 50 |
| Age (years)<br>Mean (Standard deviation) | 35 (11) | 36 (11) |
| Sex | 47% F | 44% F |
| Last year income ($)<br>Median (IQR) | 45,000 (38,000) | 35,000 (40,000) |
| Beck Depression Inventory<br>Median (IQR) | 0 (1) | 31 (13) |
| Montgomery-Asberg Depression Rating Scale<br>Median (IQR) | NA | 32 (9) |

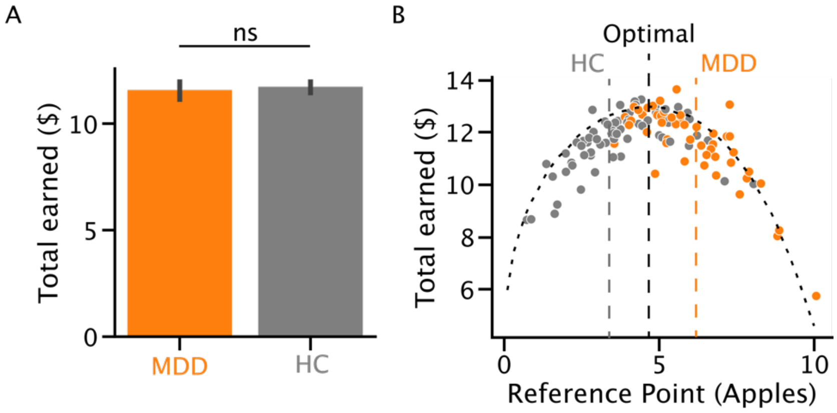


Supplementary Figure 1. Depressed participants and healthy controls had similar average amounts earned because controls tended to overharvest and depressed participants tended to underharvest. A) Average total earned in long-travel blocks during foraging session across all participants. The average earned is not significantly different between the two groups: student's t-test, $p > 0.05$. B) Total earned in long-travel blocks plotted against reference point. Total earned is optimal for a reference point of around 5 apples. However, healthy controls tended to overharvest (i.e. had a lower reference point than optimal), leading to a smaller overall average earned, while depressed participants tended to underharvest (i.e. had a higher reference point than optimal). As a result, total earned is similar across the two groups.

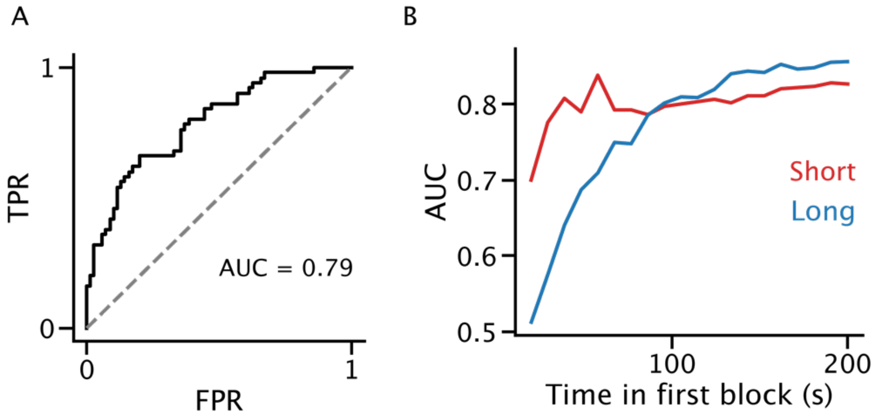


Supplementary Figure 2. Area under the curve (AUC) for short- and long-travel blocks over time. A) Receiver-operator characteristic using average short-travel reference point has an AUC of 0.79 in separating healthy control vs depressed patient. B) AUC in separating healthy control vs depressed patient for first short- or long-travel block against time in block. AUC asymptotes within around 3 minutes for both block types.

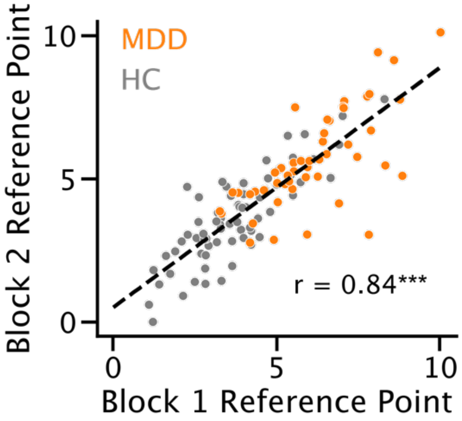


Supplementary Figure 3. Foraging task reference point is reliable between blocks. Average long-travel reference points for each participant in first and second block are highly correlated, suggesting they are reliable across the task session. Linear regression: r = 0.84, p = 1e-33.

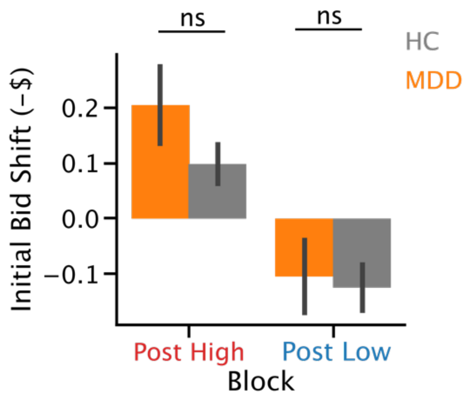


Supplementary Figure 4. Initial bidding shifts for both healthy controls and depressed participants. Average shifts in negative dollars for the first ten trials of each bidding block are not significantly different between healthy controls and depressed participants (paired-sample t-tests: $p > 0.05$ for both blocks).
